# Characterization of mutations causing steroid 21-hydroxylase deficiency in Brazilian and Portuguese populations

**DOI:** 10.1101/2021.12.07.471616

**Authors:** Mayara J. Prado, Shripriya Singh, Rodrigo Ligabue-Braun, Bruna V. Meneghetti, Thaiane Rispoli, Cristiane Kopacek, Karina Monteiro, Arnaldo Zaha, Maria L. R. Rossetti, Amit V. Pandey

## Abstract

Deficiency of Cytochrome P450 Steroid 21-hydroxylase (CYP21A2) represents 90% of cases in congenital adrenal hyperplasia (CAH), an autosomal recessive disease caused by defects in cortisol biosynthesis. Computational prediction along with functional studies are often the only way to classify variants to understand the links to disease-causing effects. Here we investigated the pathogenicity of uncharacterized variants in the CYP21A2 gene reported in the Brazilian and Portuguese populations. Physicochemical alterations, residue conservation, and effect on protein structure were accessed by computational analysis. The enzymatic performance was obtained by functional assay with the wild-type and mutant CYP21A2 proteins expressed in HEK293 cells. Computational analysis showed that p.W202R, p.E352V, and p.R484L have severely impaired the protein structure, while p.P35L, p.L199P, and p.P433L have moderate effects. The p.W202R, p.E352V, p.P433L, and p.R484L variants showed residual 21OH activity consistent with the simple virilizing phenotype. The p.P35L and p.L199P variants showed partial 21OH efficiency associated with the non-classical phenotype. Additionally, p.W202R, p.E352V and p.R484L also modified the protein expression level. We have determined how the selected CYP21A2 gene mutations affect the 21OH activity through structural and activity alteration contributing to the future diagnosis and management of 21OH deficiency.

## 1. Introduction

Congenital adrenal hyperplasia (CAH) is an autosomal recessive disease caused by defects in steroid biosynthesis [1]. More than 90% of reported CAH cases are due to 21-hydroxylase (CYP21A2) deficiency (OMIM # 201910). The CYP21A2 is a member of the cytochrome P450 superfamily and has 495 amino acids forming 13 α-helix (A-M) and 9 β-sheets [2, 3]. This protein is located in the endoplasmic reticulum of the adrenal cortex and has a role in both the glucocorticoid and mineralocorticoid biosynthesis by the hydroxylations of 17-hydroxyprogesterone into 11-deoxycortisol, and progesterone into 11-deoxycorticosterone, which are then converted into cortisol and aldosterone (Figure 1) [1]. Therefore, defects in CYP21A2 affect both the mineralocorticoid and glucocorticoid biosynthesis, besides the increase in sex steroids biosynthesis due to changes in the steroidogenesis pathway by elevated levels of sex steroid precursors [4].

**Figure 1:**
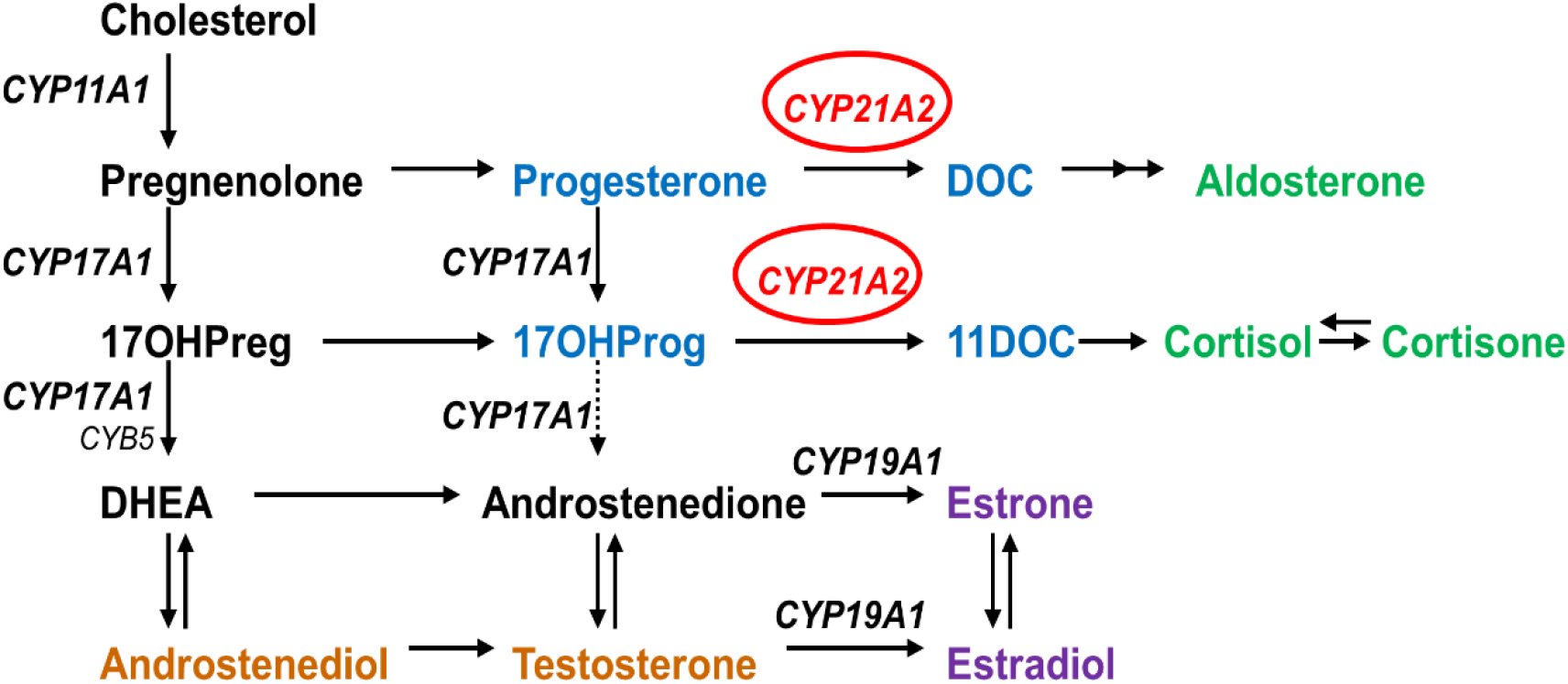
Role of CYP21A2 in human steroid biosynthesis.

Glucocorticoid and mineralocorticoid levels determine the CYP21A2 deficiency phenotype, which can be: 1) salt wasting, when both types of steroids are not produced causing the adrenal crisis by electrolytic deregulation with infant mortality risks and severe virilization by elevated sex steroids; or 2) simple virilizing, when there is some residual synthesis of both glucocorticoid and mineralocorticoid that is enough to prevent the adrenal crisis; or 3) non-classical (mild form) when just the glucocorticoid synthesis is partially affected and is linked to hyperandrogenism and mild late-onset CAH. The first two are the classical forms of CAH which have a worldwide incidence ranging from 1:14,000 to 1:18,000 live births [5]. The non-classic form has a frequency around 1:100 to 1:1000 [6]. The frequencies of CAH vary with the ethnicity and geographic population groups. However, the description of SW vs. SV, SV vs NC, etc. is rather arbitrary, hence caution must be employed when following [7]. The diagnosis of CYP21A2 deficiency is confirmed by steroid profile, mainly 17OHP in the first screening, which becomes elevated [1[5]]. However, 17OHP can be altered by a deficiency in other enzymes in the steroidogenic pathway, in premature infants, and in unrelated diseases causing physiologic stress [[8, 9]7,8]. Therefore, the molecular diagnosis is essential for the confirmation of complex cases, and follow-up management of asymptomatic CAH, avoiding unnecessary treatment, along with genetic counseling [[9, 10],[11]].

The *CYP21A2* gene is located on the short arm of chromosome 6 (6p21.3), situated 30 kb apart from its pseudogene (*CYP21A1P*). These genes share 98 % sequence identity for the exons and 96% among the introns [12]. Besides that, 95% of pathogenic variants of *CYP21A2* originate in recombination events [3]. More than 1300 variants in the CYP21A2 have been reported in the human gene mutation database (HGMD), and more than 200 of these are described to affect human health, with high variability between different ethnic groups and single nucleotide variants (SNVs), missense and nonsense account for half of the total variations [13, 14].

CYP21A2 variants are classified according to the impact on the enzyme activity. Group Null consists of deletions or nonsense variants that critically affect the enzyme activity resulting in a complete loss of function due to altered enzyme stability, steroid or heme binding, and membrane anchoring. The most common variants are: 30 kilobase deletion (30-kb del), 8 base pairs, Cluster E6 (p.I237N, p.V238E, p.M240K), p.Q319X, p.R357W and p.L307fs [3]. Group A is composed of a variant in which the enzyme activity is minimal, around 0-1%. This group is represented by an intron variant IVS2-13A/C>G which is created by an additional splice acceptor site causing retention of 19 intronic nucleotides of the intron 2 [15, 16]. Homozygous or compound heterozygote variants with the null group are often associated with the salt-wasting form, but approximately 20% of cases have a simple virilizing phenotype [3]. Group B has a residual activity of 1-10%, which is enough to prevent adrenal crisis. The p.I173N variant is representative of this group and is associated with the simple virilizing form of 21-hydroxylase deficiency [17]. Finally, group C has an enzyme activity of about 20-60% and is associated with the mild form of CAH. The variants common in this group are p.V282L, followed by p.P454S and p.P31L, which shows phenotype variability [18].

The biochemical correlation for CYP21A2 activity works well for the CAH diagnosis. However, external factors or a combination of genetic diseases can change the steroid levels, making genotype elucidation an important tool for patient management [9, 11]. In general, there is a good genotype-phenotype correlation for CAH, and the elucidation of new variants found in each population is important to improve the correct treatment [19, 20]. Initially, the bovine CYP21A2 structure was used as a template to elucidate the impact of variants damage from 2011 to 2015 (PDB ID 3QZ1), but the human CYP21A2 crystal structures have recently become available. Human CYP21A2 is deposited in the RSCB Protein Data Bank (PDB) under entries 4Y8W and 5VBU, with two different steroid ligands (progesterone in 4Y8W and 17-hydroxyprogesterone in 5VBU). The human structure has just one steroid-binding site which is different from the bovine enzyme, which has two catalytic sites [21, 22].

This study investigated the uncharacterized SNVs in the *CYP21A2* gene through computational and functional analysis, establishing a correlation with the residual enzyme activity and a possible CAH phenotype. We selected six missense variants in the *CYP21A2* gene (p.P35L, p.L199P, p.W202R, p.E352V, p.P433L, and p.R484L) that were previously reported in the Brazilian or Portuguese populations without previous functional characterization. To determine how these variants impair the CYP21A2 enzyme, we performed a detailed analysis based on the chemical changes due to amino acid alterations, residue conservation, and effect on protein structure. *In silico* predictions using human CYP21A2 structure (PDB ID 4y8w) were used to understand the structural damage, while the *in vitro* analysis using recombinant protein expression and enzyme kinetics was performed to determine the impact on protein function.

## 2. Results

### 2.1. Variant collection and description

We selected six missense variants that had patient genotype and phenotype available in our literature review after the initial screening described in methods. Four of these variants were found in Brazilian (p.P35L, p.L199P, p.E352V, and p.R484L) and two in Portuguese (p.W202R and p.P433L) populations. Genetic and clinical data describing the carriers of the selected SNVs are summarized in **Table 1**. These data were collected from the original papers.

**Table 1.** Genetic and clinical features of the subjects carrying the selected SNVs (bold) in the *CYP21A2* gene. All data are from the original papers. Normal values for 17α-hydroxyprogesterone (17OHP) is < 30 nmol/L, serum sodium between 132-142 mmol/L and potassium 3.6-6.1 mmol/L. LGC, large gene conversion; 30-kb del, large deletion from *CYP21A1P* to *CYP21A2* gene; nd, non-determined; DSD, ambiguous/atypical genitalia; AC, adrenal crisis; M, male; F, female. ^a^ Age at diagnosis. ^b^ non-diluted measurements (reference value 5.97 nmol/L). ^c^ after ACTH stimulation.

| Genotypes |  | Phenotype | Clinical data | Sex | 17OHP (nmol/L) | Na<sup>+</sup>/K<sup>+</sup> (mmol/L) | Age <sup>a</sup> | Reference |
| --- | --- | --- | --- | --- | --- | --- | --- | --- |
| Allele 1 | Allele 2 |  |  |  |  |  |  |  |
| **P35L**, H63L, 30-kb del | LGC | SW | vomiting, dehydration | M | 49.3 | 116/9.2 | 2.8 m | [23] Coeli et al. (2010) |
| **P35L**, H63L, 30-kb del | Q319X | SW | vomiting, dehydration | M | > 25 <sup>b</sup> | 120/6.8 | 13 m | [23] Coeli et al. (2010) |
| **P35L**, H63L, 30-kb del | c.920_921insT | SW | DSD (Prader IV) | F | > 6 <sup>b</sup> | 119/5.3 | 15 d | [23] Coeli et al. (2010) |
| **P35L**, H63L, 30-kb del | R357W | SW | vomiting, dehydration, AC | M | > 200 <sup>b</sup> | 119/9.7 | 28 d | [23] Coeli et al. (2010) |
| **L199P** | normal | ND | DSD (Prader I) | F | 408.5 | nd |  | [24] Silveira et al. (2009) |
| **W202R** | LGC | SW | DSD, salt wasting | F | nd | nd | < 2 m | [25] Santos-Silva et al. (2019) |
| **E352V** | E6 cluster | SV | nd | F | nd | nd | nd | [10] De Carvalho et al. (2016) |
| **E352V** | IVS-13A/C>G | SW | nd | M | nd | nd | nd | [10] De Carvalho et al. (2016) |
| **E352V** | IVS-13A/C>G | SW | nd | M | nd | nd | nd | [10] De Carvalho et al. (2016) |
| **E352V** | G425S | SV | nd | M | nd | nd | nd | [10] De Carvalho et al. (2016) |
| **P433L** | P454S | NC | nd | F | 42.7 <sup>c</sup> | nd | 17 y | [26] Carvalho et al. (2012) |
| **R484L**, IVS-13A/C>G | Del CYP21A2 | SW | DSD (Prader III-IV) | F | 1537.2 | 134/4.9 | < 4 m | [24] Silveira et al. (2009) |
| **R484L**, IVS-13A/C>G | Q319X | SW | undefined | M | 1358.7 | 130/6.2 | < 9 m | [24] Silveira et al. (2009) |

p.P35L was identified in four Brazilian patients as compound heterozygous with other mutations (**Table 1**). One allele presents a mutation with symptoms of severe enzyme damage, while the other alleles carried three mutations, p.P35L, p.H63L and 30-kb del. It was concluded that p.P35L origin was not from the same genetic event of the p.H63L and 30-kb del, as it was not present in the pseudogene screening. Therefore, we analyzed this variant alone.

p.L199P was found as heterozygous in a female patient with mild clitoromegaly atypical genitalia and elevated 17OHP level detected in the Brazilian newborn screening program (**Table 1**). According to the authors, the child remained asymptomatic at 3.3 years old, and probably the clitoromegaly and the high 17OHP level detected in the screening were due to the premature birth. The impact of this new variant has been unknown as it was not possible to infer the consequence once it was identified in heterozygosis.

p.W202R was found as compound heterozygous with large gene conversion in a Portuguese female (**Table 1**). This patient presented atypical genitalia and salt wasting, being diagnosed with classical CAH at < 2-month-old.

p.E352V was identified in a compound heterozygous state in four classical CAH patients from Brazil (**Table 1**). Two of them with the same intronic mutation IVS2-13A/C>G heterozygous with p.E52V. These two patients showed the classical CAH form with salt-wasting. One female presented p.R484L in compound heterozygous with Cluster E6. These patients presented the simple virilizing CAH form. The latter has the p.R484L and p.G425S which resulted in the simple virilization form of CAH.

p.P433L was identified in a Portuguese female who presented it in compound heterozygous with the mild mutation p.P454S (**Table 1**). This patient was diagnosed at 17-years-old with a nonclassical form of CAH after ACTH stimulation, which indicated the 17OHP elevation characteristic of that form of CAH.

p.R484L was found as compound heterozygous in two Brazilian patients (**Table 1**). Both present classical CAH form, with salt-wasting and high 17OHP levels detected during newborn screening. One allele in both cases has the p.R484L together with a splice variant IVS2-13A/C>G. The second allele in one patient was a CYP21A2 deletion and in the other p.Q319X.

### 2.2 Computational characterization indicated the structural impact of the SNVs

We performed a screening with five predictive tools that have different approaches, PolyPhen-2, SNAP2, MutPred2, Meta-SNP, and PredictSNP. Results obtained for the six variants chosen in this work are shown in Table 2. p.L199P, p.E352V, and p.R484L were predicted to damage the protein by all predictor tools, while p.P35L had damage predicted by 4 tools, p.W202R by 3 and p.P433L by 2 different tools.

**Table 2.** Prediction of the possible impact on the structure and/or function of CYP21A2 by genetic variants. Seven variants were used to validate the method, with four polymorphisms (●) and three with a known negative impact (↓). ^a^ SNV nomenclature according to UniProt ID Q16874-1. Scores ≥ 0.5 by PolyPhen-2, MutPred2, and Meta-SNP indicate protein damage. SNAP2 scores >50 indicate a strong signal for effect, between 50 and −50 weak signal and <-50 strong signals for neutral effect.

| SNV<sup>a</sup> | *In silico tools* |  |  |  |  |  |  |  |  |
| --- | --- | --- | --- | --- | --- | --- | --- | --- | --- |
|  | PolyPhen-2 | Score | SNAP2 | Score | MutPred2 | Score | Meta-SNP | Score | PredictSNP |
| ● R103L | Benign | 0.023 | Neutral | -47 | Neutral | 0.191 | Neutral | 0.310 | Neutral |
| ● D184E | Benign | 0.000 | Neutral | -85 | Neutral | 0.260 | Neutral | 0.412 | Neutral |
| ● S269T | Benign | 0.016 | Neutral | -93 | Neutral | 0.076 | Neutral | 0.340 | Neutral |
| ↓ I173N | Damage | 1.000 | Effect | 70 | Pathogenic | 0.838 | Disease | 0.811 | Deleterious |
| ↓ V282L | Benign | 0.273 | Neutral | -76 | Neutral | 0.084 | Neutral | 0.344 | Neutral |
| ↓ R427H | Damage | 1.000 | Effect | 82 | Pathogenic | 0.821 | Disease | 0.827 | Deleterious |
| P35L | Damage | 0.988 | Effect | 13 | Pathogenic | 0.676 | Disease | 0.624 | Neutral |
| L199P | Damage | 0.997 | Effect | 68 | Pathogenic | 0.875 | Disease | 0.711 | Deleterious |
| W202R | Damage | 1.000 | Effect | 45 | Pathogenic | 0.577 | Neutral | 0.451 | Neutral |
| E352V | Damage | 0.693 | Effect | 90 | Pathogenic | 0.919 | Disease | 0.937 | Deleterious |
| P433L | Benign | 0.402 | Effect | 15 | Pathogenic | 0.614 | Neutral | 0.372 | Neutral |
| R484L | Damage | 1.000 | Effect | 81 | Pathogenic | 0.690 | Disease | 0.748 | Deleterious |

Among the variants with known activity, the neutral variants (p.R103K, p.D183E, and p.S269T) agreed with all tools. However, among the three variants known to cause damage, we had different results. The two severe variants (p.I173N and p.R427H) were predicted correctly by all tools, however, the mild (p.V282L) variant was wrongly classified by all of them. The second screening about the protein stability of all variants presented a variation of ΔG, from −1.47 kcal/mol (p.R484L) to 1.23 kcal/mol (p.E352V) (**Table 2 and 3**).

**Table 3.** Summary of structural and functional data obtained for the six variants studied shows that all of them can be related to CYP21A2deficiency. ^a^ SNV nomenclature according to UniProt ID Q16874-1. ^b^ two hydrogen bonds with E355. H, hydrogen-bound.

| SNV <sup>a</sup> | Structure/protein localization | Physicochemical properties | | $\Delta\Delta G$<br>(kcal/mol) | Enzyme activity<br>(Mean ± SD) | The molecular mechanism<br>(Hypothesis) | Mutation group |
| --- | --- | --- | --- | --- | --- | --- | --- |
|  |  | WT | MUT |  |  |  |  |
| **P35L** | N-term coil / Region of the membrane protein orientation | Polar residue with a ring at the N-term | Hydrophobic; Short side chain; Folding interactions; | -0.58 | 13.2 ± 1.56% | Disturbs the protein orientation in the membrane | C |
| **L199P** | F-helix / Close to the active site; α-helix stabilization | Hydrophobic; Short side chain; Folding Interactions; H: I195 and S203 | Polar residue with a ring at the N-term; H: S203 | 0.79 | 10.3 ± 0.33% | Breaks the secondary structure close to the active site | C |
| **W202R** | Turn of the F-helix / close to the active site and steroid bound residue (R234) | Hydrophobic; Large, rigid aromatic group; H: V198 | Hydrophobic; Long, flexible, and posit. charged side chain; Ionics bound; H: V198 | 1.11 | 1.2 ± 0.34% | Positive charge addition causes disorder in heme binding | B |
| **E352V** | K-helix / ERR-triad associated with the heme group | Hydrophobic; Long, slightly, flexible side chain; Strongly neg. charged; Ionic bound; Fix metal iron; H: A348, R355<sup>b</sup>, L356 and W406 | Hydrophobic; Short side chain; Folding interactions; H: A348 and L356 | 1.23 | 1.1 ± 0.11% | Disorder of the ERR-triad decreases heme-binding and stability | B |
| **P433L** | L-helix/ Adjacent to essential residues for the heme bond | Polar residue with a ring at the N-term; H: A435 | Hydrophobic; Short side chain; Folding interactions; H: A435 | 0.39 | 7.5 ± 0.67% | Changes the natural orientation of heme-binding residues R427 and C429 | B |
| **R484L** | C-term coil / Hydrophobic cluster with ionic connection | Hydrophobic; Long, flexible, and posit. charged side chain; Ionics bound; H: Q482, A449 and M486 | Hydrophobic; Short side chain; Folding interactions; H: A449 | -1.47 | 3.4 ± 0.8% | Loss of hydrophobic organization at the C-term region | B |

### 2.3 Amino acid chemical proprieties, conservation, and structural damage

We analyzed the amino acid chemical properties to assess the characteristics of each exchange, and the conservation of that residue across species to determine how it has been retained during the evolution of CYPs P450. We worked with 200 CYP 450 sequences homologous to human CYP21A2. The structural damage was characterized by Gibbs’s free energy and gain/loss of hydrogen bonds.

Proline at amino acid position 35 is located in a coil on the protein surface (Figure 2). This residue shows a high conservation score, with only two residues found on homologous sequences (proline and guanine) (Figure 4 and Table 4). p.P35L has the amino acid properties changed from a polar and uncharged to a nonpolar and aliphatic residue, increasing the structural stability, ΔΔG −0.58 kcal/mol (Table 3). PolyPhen-2, SNAP, Meta-SNP, and MutPred2 predicted the effect of this mutation on the protein stability and functionality. Besides that, MutPred2 predicts two structural effects: gain of helix and alteration of transmembrane features. PredictSNP classified this variant as neutral (Table 2).

**Table 4.** ConSurf amino acid conservation score. The score is from 1 (variable) to 9 (conserved). CI - Confidence interval of the score. ^a^ PDB ID 4Y8W.

| SNV <sup>a</sup> | Conservation score | CI | Residue variety |
| --- | --- | --- | --- |
| **P35** | 9 | 9 to 8 | L, P, G |
| **L199** | 3 | 4 to 2 | T, N, L, V, F, I, A, S, G, M |
| **W202** | 1 | 2 to 1 | T, E, R, W, G, C, M, L, A, I, N, Q, D, V, F, S |
| **E352** | 9 | 9 to 9 | E |
| **P433** | 3 | 4 to 2 | R, K, G, M, L, V, H, A, S, I, T, P, E, Q |
| **R484** | 9 | 9 to 9 | R, T, L |

**Figure 2.**
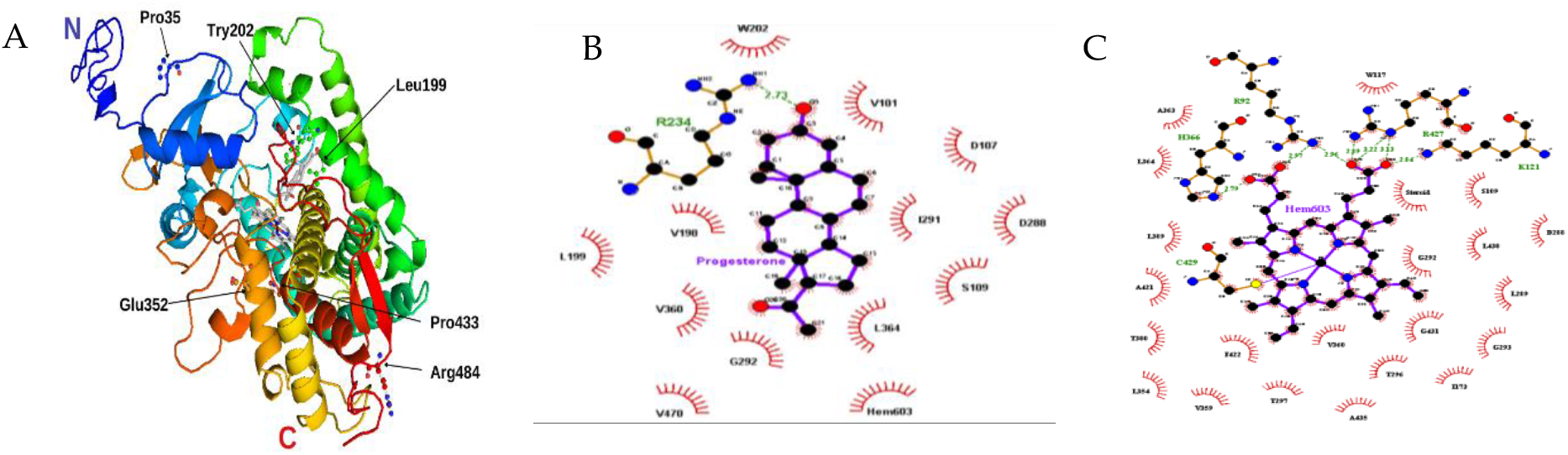
Structural model of CYP21A2. **(A)** We built the missing extremity based on PDB ID 4Y8W with I-TASSER and visualized by PyMOL. The amino acids mutated in our study are identified with black arrows. Progesterone and protoporphyrin containing Fe^3+^ are in the middle of the structure colored in grey **(B)** Contact map with progesterone binding site in 21OH structure. The studied residue p.W202 is shown on the top. **(C)** Contact map with heme (protoporphyrin containing Fe) binding site in 21OH structure. The residue p.C429 and p.R427 are near the studied residue p.P433. Heme and progesterone contact maps were built with the structure of CYP21A2 (PDB # 4Y8W). Residue ligand contact map were generated with LigPlot+ v.2.2.4 software using as maximum hydrogen -acceptor / -donor distance 2.7 Å and 3.35 Å (green line), respectively, while for non-bonded, the minimum contact distance was 2.9 Å and the maximum 3.9 Å.

Leucine at position 199 is localized at α-helix F, close to the CYP21A2 catalytic site < 4 Å from heme, where it forms two hydrogen-bonds, one with p.S203 (on the loop between α-helix F and F’) and one with p.I195, on α-helix F (Figure 3 and Table S1). This residue shows mild conservation, being found in 10 different sequences at this position (Figure 4, Table 4). p.L199P changes a nonpolar and aliphatic residue to a polar and uncharged residue, losing the hydrogen bond with p.I195 (Table S2). All predictor tools showed that this mutation causes damage to the protein, and MutPred2 predicted an alteration of coiled-coil and transmembrane regions. ΔΔG was 0.79 kcal/mol.

**Figure 3.**
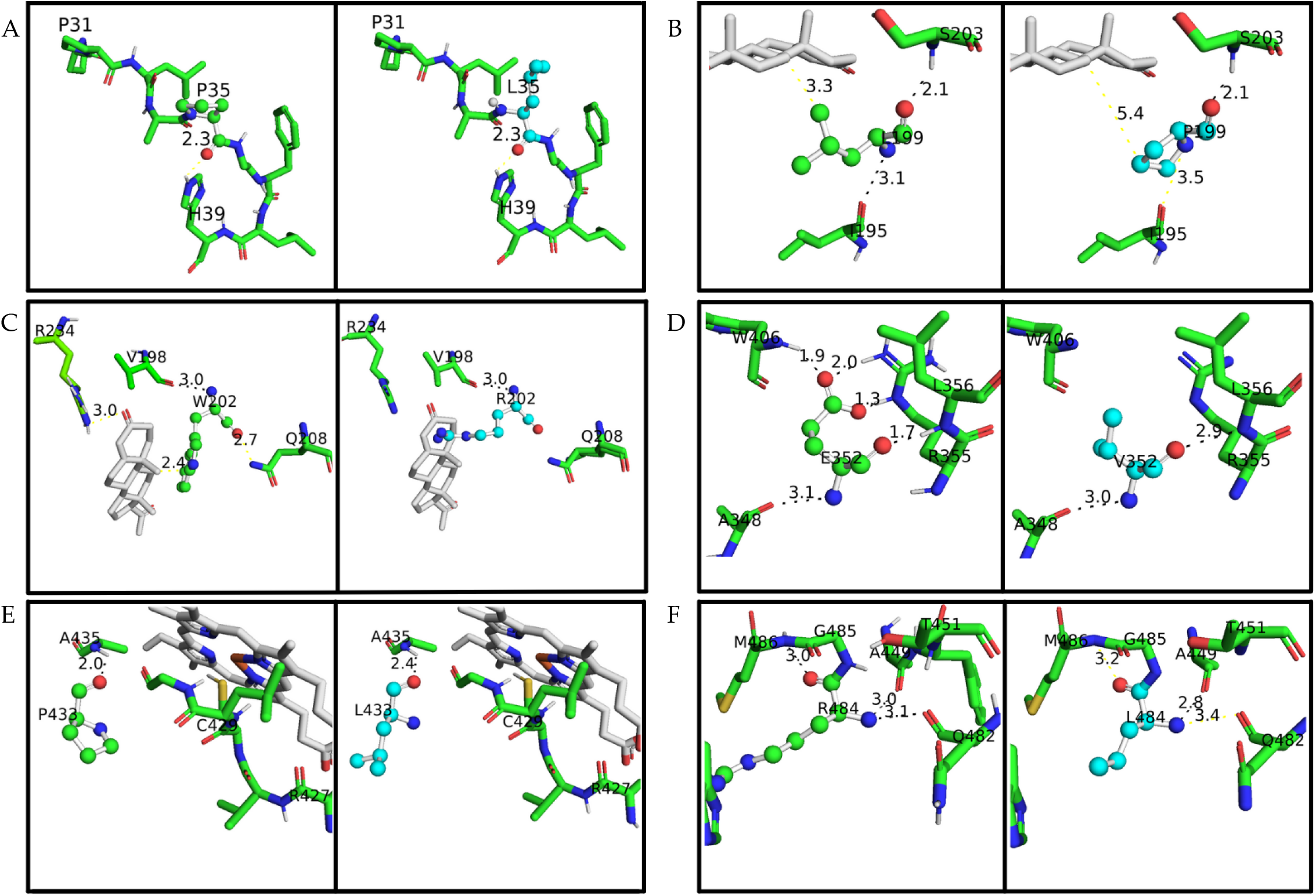
A closeup of the amino acid studied on the CYP21A2 structure. The protein structure is based on PDB ID 4Y8W. Wild-type amino acids are colored in green and mutated in cyan, in both of them oxygen is red, nitrogen is blue, and sulfur is yellow. The distance measured between the main amino acids connections is with less than 4 Å. Hydrogen bonds are represented with black lines, while any other measurement between atoms is with yellow lines. Progesterone and heme are colored grey. (A) p.P35 and p.P35L, (B) p.L199 and p.L199P, (C) p.W202 and p.W202R, (D) p.E352 and p.E352V, (E) p.P433 and p.P433L, (F) p.R484 and p.R484L.

**Figure 4.**
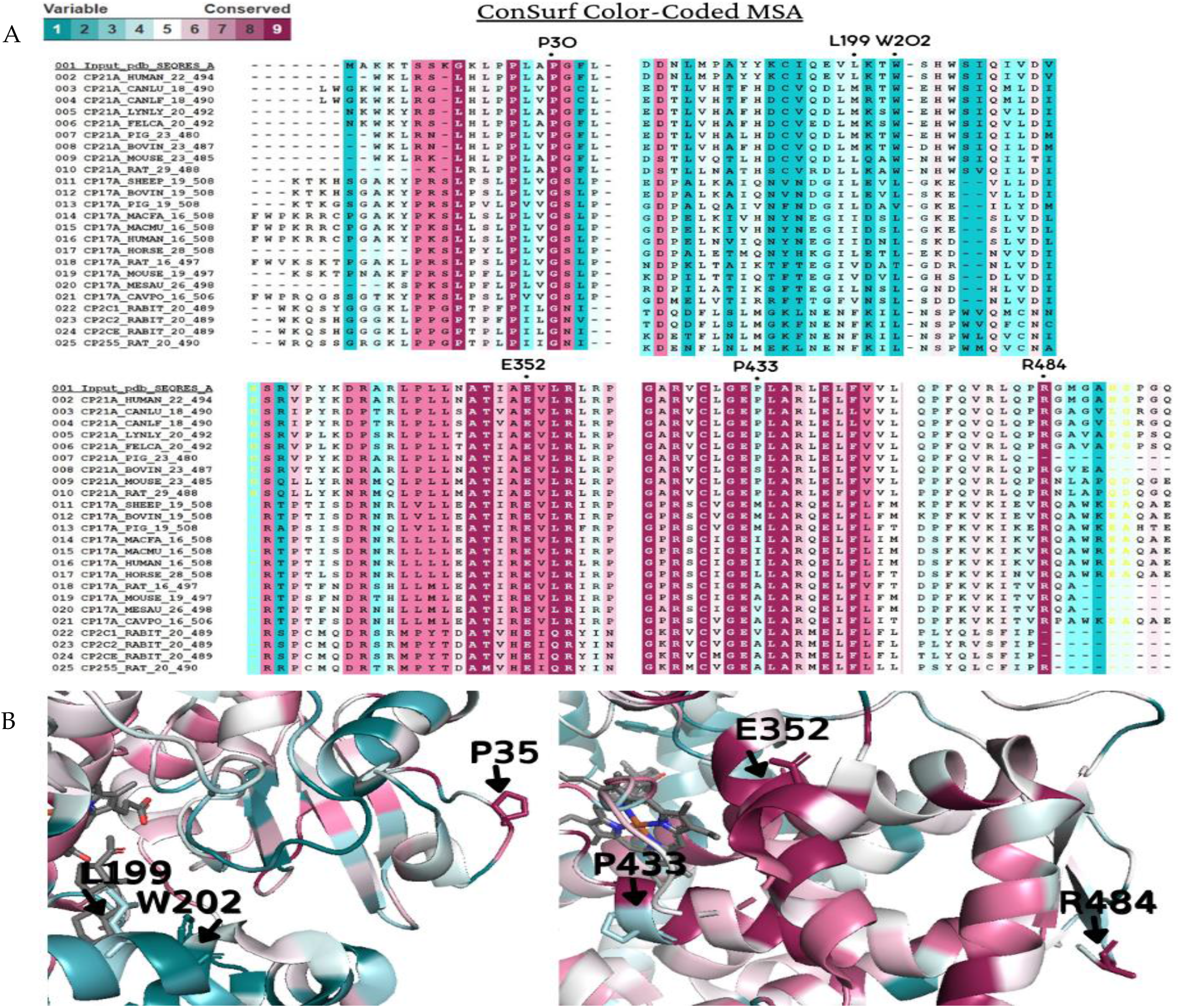
Amino acid conservation of the human CYP21A2.: A CYP21A2 structure (PDB # 4Y8W) structure was used as a reference for ConSurf analysis. Multiple sequence alignment was built using CLUSTALW with 200-cytochrome P450 sequences homologous to the human protein 21-hydroxylase. The species considered for final alignment were from *Homo sapiens, Bos taurus, Canis lupus familiaris, Cavia porcellus, Capra hircus, Felis catus, Gorilla gorilla gorilla, Lynx lynx, Mesocricetus auratus, Macaca fascicularis, Macaca mulatta, Mus musculus, Oryctolagus cuniculus, Ovis aries, Rattus norvegicus* and *Sus scrofa*. **(A)** A representative part of the alignment from MSA. The specific positions of the six wild-type residues analyzed in this work are marks with a dot (●). **(B**) CYP21A2 structure with black marks on the six wild-types residues on the protein structure. Residues p.P35, p.E352, and p.R484 showed to be conserved, while p.L199, p.W202, and p.P433 are variable. Color-code: from pink (conserved) to cyan (variable).

Tryptophan at position 202 is also located < 4 Å from the CYP21A2 catalytic site, on the α-helix F with a hydrogen bond with p.V198 (Figure 3). This residue presented a low conservation score, being highly variable between its homologous sequences (Figure 4 and Table 4). p.W202R changes a hydrophobic residue to a hydrophilic and positively charged residue, decreasing the CYP21A2 structural stability with a ΔΔG of 1.11 kcal/mol, but without losing the hydrogen bond (Table S2). Two predictor tools classified that residue replacement as neutral (Meta-SNP and PredictSNP), while the three others predicted damage to the protein. An altered coiled-coil region was predicted by MutPred2.

Glutamate at position 352 is located in α-helix K, presenting five hydrogen bonds, four on the same helix (one with p.A348 and p.L356, and two with p.R355)) and one with a coil (p.W406) (Figure 3 and Table S1). This residue is highly conserved across species, and there is no other variation on CYP21A2 homologs (Figure 4 and Table 4). P.E352V changes a negatively charged residue to a nonpolar aliphatic residue. The stability of the CYP21A2 structure was increased, and the changed residue lost three hydrogen bonds with W406, and p.R355 (Table S2). All the predictor tools showed that p.E352V results in protein damage (Table 2). The MutPred2 predicted alteration of an interface, loss of allosteric site at p.E352, and altered metal binding.

Proline at position 433 is located at α-helix M, where it has a hydrogen bond with p.A435 in the same helix (Figure 3 and Table S1). This residue is close to the central heme group, but it is not a conserved residue, as it is found interchangeable as 14 different residues among the CYP21A2 homologous group (Figure 4 and Table 4). However, this indicates the evolution of this residue across species with different roles and different redox partners. The exchange of a polar and uncharged amino acid with a nonpolar and aliphatic showed ΔΔG 0.39 kcal/mol (Table 3). SNAP2 predictor tool showed damage on the protein with the variant, and MutPred2 a gain of helix and loss of catalytic site at p.E432, however, the other three predictors indicated a neutral effect (Table 2).

Arginine at position 484 is located on the protein surface in a coil which makes a cluster with 9 other amino acids. This residue has three hydrogen bonds, one with p.M486 in the same coil, one with p.Q482, located at stand β9, and one with p.A449, which makes the connection between α-helix M and β8 (Figure 3 and Table S1). This residue showed the highest conservation score, with just two other amino acids found in the homology analysis (Figure 4 and Table 4). The variant p.R484L changes a positively charged polar residue to a nonpolar and uncharged residue, losing the two hydrogen bonds with p.M486, and Q482 (Table S2). This amino acid exchange showed damage by all predictor tools used and stability increased with a ΔΔG of −1.47 kcal/mol (Table 2 and Table 3). Loss of intrinsic disorder was predicted by MutPred2 as well.

### 2.4 Functional testing for the validation of in silico results

HEK293 cells were transfected with plasmids expressing CYP21A2 WT or variants p.P35L, p.L199P, p.W202R, p.E352V, p.P433L, and p.R484L, and p.I173N and p.V282L as two controls with known activity (∼2 % and 18-60 %, respectively) [4, 17, 18]. The CYP21A2 activity was quantified for both WT and variants. Using the TLC analysis, we could assess the conversion ratio of progesterone to 11-deoxycorticosterone and compare the activity of each variant with the WT enzyme (Figure 5A). Variants p.W202R and p.E352V showed only residual conversion (< 2% enzyme activity), similar with p.I73N, which is associated with the classical form of CAH. Variants p.P35L and p.L199P showed partial activity, with the catalytic activities 13.2% and 10.3% of the WT, similar with the p.V282L variant, which is associated with a nonclassical form of CAH. Variants p.P433L and p.R484L presented activity between the two controls, being 7.5% and 3.4% of the WT activity respectively (Figure 5C).

**Figure 5.**
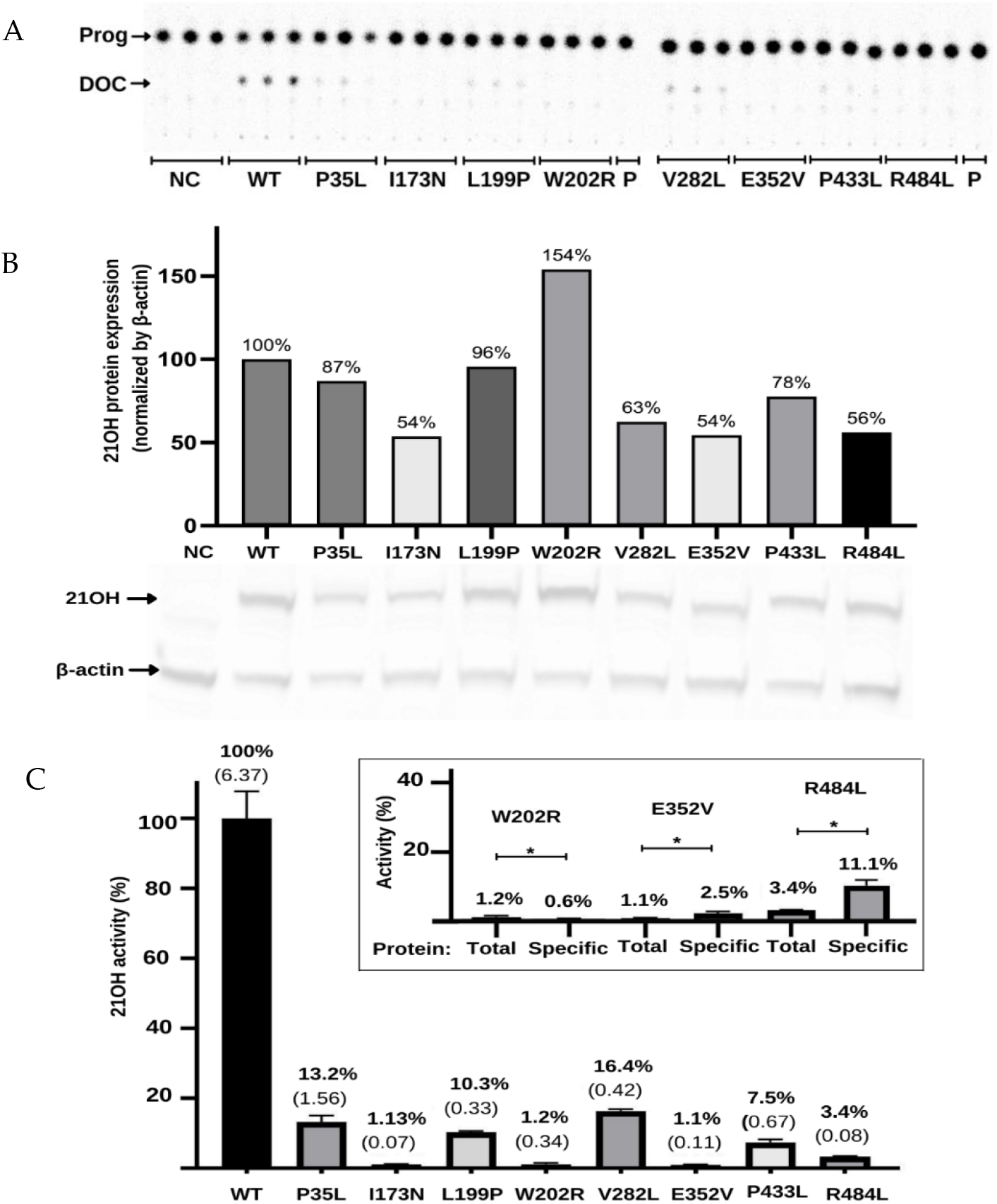
Functional characterization of the variants in the *CYP21A2* gene by relative steroid conversion of progesterone (Prog; P) to 11-deoxyprogesterone (DOC). The activity was obtained using 21OH wild-type (WT) and mutated expressed in HEK293t cell where we measured the steroid conversion by the percentage of radioactivity into DOC to the whole sample (A) Thin-layer chromatography (TLC) shows that all variants reduce the activity of 21-hydroxylase (21OH). **(B)** Semiquantitative 21OH expression level obtained by western blot with anti-flag antibody for the 21OH (53 KDa) and normalized by β-actin (42 KDa) expression with anti-β-actin antibody. Negative control (NC) presents the basal HEK293 cells. **(C)** 21OH activity expressed accordingly with TLC spots densitometry and related to the WT. Bars represent the standard error from three samples. **Inside the box** is plotted the activity (total) and the specific activity (specific) of the three variants that showed statistic difference (p<0.05) between these values. The specific activity was obtained by dividing the enzyme activity by the 21OH protein estimated by western blot. All results were analyzed on GraphPad software.

To determine if the reduction of activity was associated with the decrease in protein expression, we quantified CYP21A2 protein in WT and variants using western blot assay (Figure 5B). Normalized by β-actin expression, we found the expression level was less than 50% of the WT for the variants p.P35L, p.V282L, p.E352V, and p.R484L variants, while p.W202R had 119% of WT. Therefore, we also calculated the specific activity of each variant compared to WT by dividing the activity obtained from the TLC results by the CYP21A2 expression level (Figure 5C). The specificity activity for p.W202R, p.E352V, and p.R484L present a significant difference (p < 0.05) when compared with the non-protein normalized activity. p.W202R had a decrease of activity to < 1%, while p.E352V and p.R484L had an increased activity by 2 and 3-fold, respectively. Altogether, our results indicated that all variants tested significantly impacted the activity of CYP21A2. The variants p.W202R, p.E352V, and p.R484L also impacted the protein expression (Figure 5C).

### 2.5 Kinetic analysis of CYP21A2 variants

The apparent kinetic constant revelated saturation for the 21OH WT with the Michaelis-Menten constant (K_m_) of 1.57 µM for progesterone and an apparent maximal reaction velocity (V_max_) of 0.360 nmol.min-1.mg-1 (**Figure 6 and Table 4**). The apparent saturation was a similar rate of the WT for p.P35L (K_m_ 2.06 µM), and p.P433L (Km 1.91 µM), but it was six times higher for p.L199P (K_m_ 10.24 µM), indicating that p.L199P decreases the protein affinity for the substrate progesterone. The apparent V_max_ was lower than the WT for p.Pro35Leu (0.085 nmol.min^-1^.mg^-1^), p.L199P (0.252 nmol.min^-1^.mg^-1^) and p.P433L (0.055 nmol.min^-1^.mg^-1^). The apparent catalytic efficiencies of the mutated proteins were also lower than those of WT, p.P35L had catalytic efficiency of 18% compared to the WT, p.L199P of 11%, and p.P433L of 13% (**Table 5, Figure 6**).

**Table 5.** Apparent kinetic constants and catalytic efficiency were calculated from 3 independent experiments.

|  | Wild<br>type | P35L | L199P | P433L |
| --- | --- | --- | --- | --- |
| $V_{\text{max}}$ (nmol.min<sup>-1</sup>.mg<sup>-1</sup>) | 0.360 | 0.086 | 0.252 | 0.055 |
| $K_{\text{m}}$ ($\mu\text{M}$) | 1.57 | 2.07 | 10.24 | 1.91 |
| $V_{\text{max}}/K_{\text{m}}$ | 0.229 | 0.041 | 0.025 | 0.029 |

**Figure 6.**
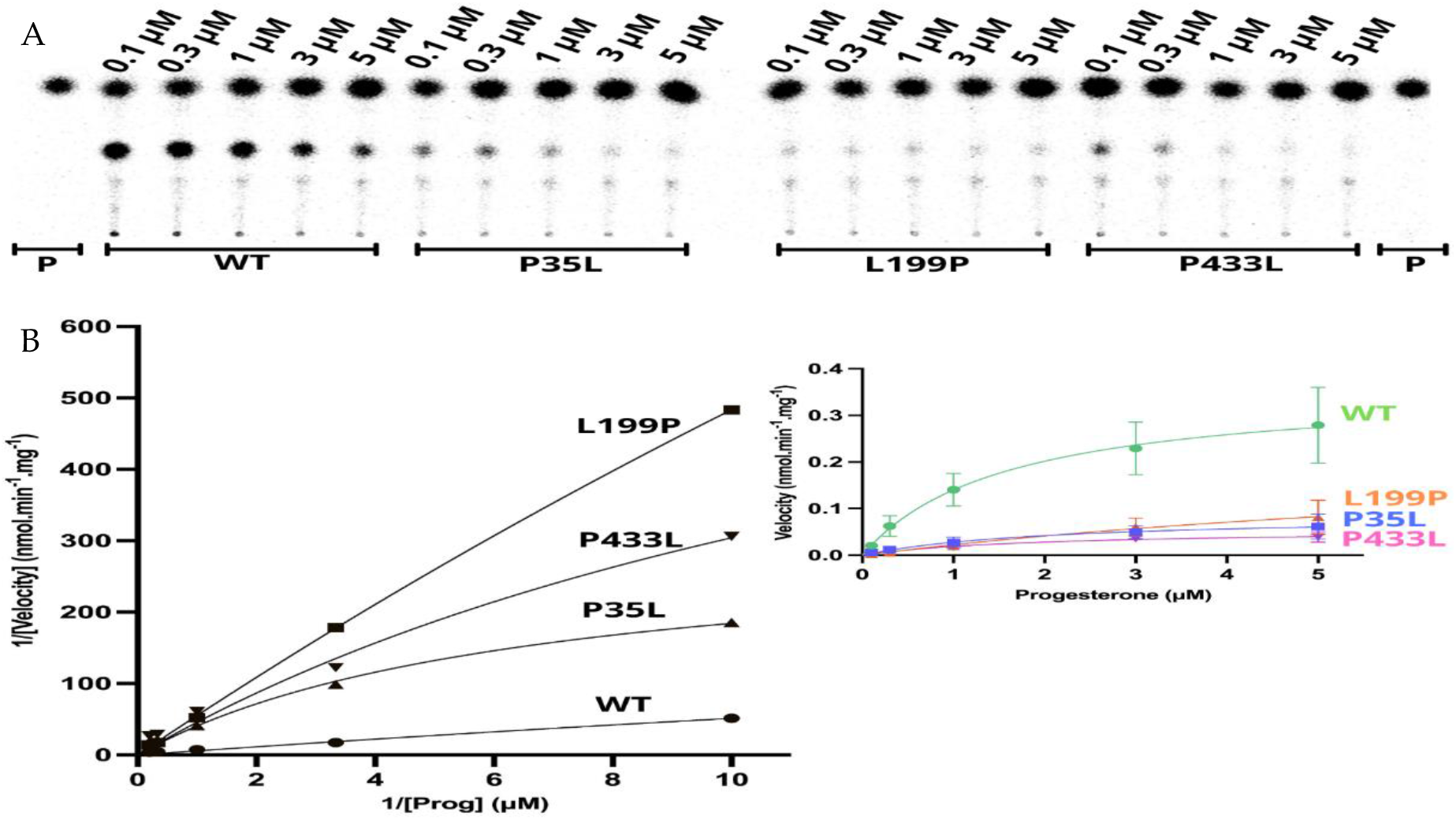
Kinetics assay shows a decrease of reaction velocity for three variants which have still presented activity in the functional assay. (**A**) representative thin layer chromatography (TLC) from three replicates shows the CYP21A2 conversion of progesterone (Prog) (0.1, 0.3, 1, 3, and 5 µM) into 11-deoxycorticosterone (DOC). (**B**) **The left-hand plot** presents the linear plots of enzymatic activity of 21OH WT and mutated between 1/Velocity against 1/Prog (progesterone concentration) for the conversion of progesterone to DOC. Additionally, the **right-hand plot** presents the Michaelis Menten nonlinear curve with velocity reactions against the substrate progesterone used to obtain V_max_ and K_m_ values.

Our results show that the p.L199P variant, which is located close to the catalytic site, affects the substrate binding. In contrast, p.P35L and p.P433L showed inhibition of the enzyme activity through a decrease in reaction velocity.

## 3. Discussion

We have presented a detailed investigation of the molecular and functional damage caused by six missense variants of the *CYP21A2* gene. These variants were selected from a literature review of uncharacterized missense variants. We performed *in silico* analysis as well as functional assays to assess the effect of these CYP21A2 variants. We showed the analysis of two variants with changes in positions close to the catalytic site, p.L199P, and p.W202R. Both showed a low conservation score among mammalian CYPs P450. However, looking specifically for the CYP21A2, residue 199 still had variability while 202 had high conservation. The p.W202R changed a heavy and hydrophobic residue to a hydrophilic and positively charged residue. This exchange at 202 abolished the enzyme activity, which is in agreement with the severe CAH (SW) form described for the patient [25]. In the other case, the leucine replacement by proline at position 199 broke the hydrogen bond with a residue on the same helix, destabilizing the conformation of helix F. Previous studies with helical substitution of leucine to proline at position 168 (helix E), 262 (helix H), and 322 (helix J) showed a break of the helix as well, and association with the severe form of CAH [4, 19, 27]. In our study, this alteration decreased the affinity of the enzyme for the substrate by about 6-fold, making the enzyme activity drop to 10 % of the WT.

We analyzed two CYP21A2 variants by looking at the protein surface, one at the N-terminal, residue 35, and the other on the C-terminal, residue 484. We showed that the proline at the 35 position has a high conservation score, even being on the protein surface without strong interaction with closer residues. This variant decreases the reaction velocity 4 times due to the faster enzyme saturation, V_max_. 0.08552 nmol.min^-1^.mg^-1^, compared with WT, V_max_. 0.3604 nmol.min^-1^.mg^-1^. The apparent enzyme activity was around 34.8 % of the WT. That result suggests an important and conserved role of P35 across species. Variant p.P35L was previously described in the Brazilian population in a compound heterozygous state with other three variants, being two in the same allele and one in the other [23]. All the four patients described by [23] carry two other pathogenic mutations (30-kb del and p.H63L) at the same allele of p.P35L, which by themselves would transcribe a non-functional protein [3, 28]. The p.H63L is derived from pseudogene and has known mild activity in the homozygous state, however, it has a synergistic effect when it is associated with another mild mutation and results in a severe phenotype [28]. The 30-kb del is a gene rearrangement event on RP1-C4A-CYP21A1P-TNXA and RP2-C4B-CYP21A2-TNXB region, which causes CYP21A2/CYP21A1P chimera [3]. In [23], p.P35L was not presented on any pseudogene analyzed, which indicated that these appear in the same allele from independent genetics events.

The residue studied at the carboxyl-terminus, 484, has high conservation while all the others around this sequence position have mild or low conservation scores. The p.R484 stabilizes other residues through hydrogen bonds with p.M486 in a coil, p.Q482 at stand β9, and p.A449, a crucial residue between helix M and a sheet β8 [29]. The variant p.R484L changes a polar and positively charged amino acid to a nonpolar and uncharged residue inside a polar cluster, causing an intrinsic disorder. This variant showed enzyme activity around 11% of the WT when normalized by the specific amount of CYP21A2. However, the activity was significantly lower without the protein normalization, suggesting this variant generates a lower transcription level or higher protein degradation than the WT. The same effect was observed for p.W202R and p.E352V variants. The p.R484L was found in a compound heterozygous state with other two variants, one in the same allele and another on the opposite allele [24] both of these have known severe impact on enzyme activity [3, 20].

Another residue with a high interaction number is the p.E352. This amino acid is highly conserved, and there is no residue variation at the same position in homologous sequences, showing an important and conserved role in the CYP21A2 protein. The p.E352 forms five hydrogen bonds, stabilizing its helix through four interactions (two with p.A348 and p.L356, and two with p.R355), and stabilizing a network with p.W406, in the L-M loop. The tryptophan at 406 is a backbone residue, while arginine at 355 has an extensive network with glutamine at p.E352 and arginine at 409 in the ERR-triad [19]. This triad acts to stabilize the three-dimensional structure that allows covalent binding of the heme group [19, 30]. The variant p.E352V causes the loss of three hydrogen bonds (p.W406, and p.R355), destabilizing several structural elements. We showed that the CYP21A2 p.E352V decreased the enzyme activity to only 2.5% of the WT, which would be related to the classical form of CAH. de Carvalho et al [10] identified four patients with this variant in the compound heterozygous state, two of them had a simple virilization form of CAH, while two had the salt-wasting form. In the first form, one patient presented Cluster E6, which abolishes the enzyme activity, and the other p.G425S, which has less than 2% of enzyme activity [4]. The latter form was detected in two patients with the same intronic variant IVS2-13A/C>G, which mostly results in the salt-wasting form (in around 79% of the cases) and simple virilizing form (in 20% of the cases) [3].

Lastly, the residue p.P433 had a low conservation score looking at the mammalian CYPs P450. Nonetheless, the score was high among the CYP21 group, highlighting the importance and conservation of this residue in this specific group. Proline at 433 is on the first helix M turn and it has one hydrogen bond with a residue in the same helix M, p.A435, helping with the helix stabilization. Besides that, this residue lies adjacent to the L-M loop, where a heme-binding residue p.R427 is located. The p.P433L makes the structure more flexible, modifying the optimal conditions for the heme-binding, as previously described [19]. This variant makes the reaction become saturated 6 times faster than the WT, with a V_max_. of 0.055 nmol.min^-1^.mg^-1^, and decreased the enzyme activity to ∼11.4 %. The p.P433L was identified in the compound heterozygous state with the mild mutation p.P454S in a patient diagnosed with the nonclassical form of CAH [26].

## 4. Materials and Methods

### 4.1 Search and selection of variants in CYP21A2

We searched Ensembl, HGMD, OMIM, and ClinVar databases for all *CYP21A2* variants with an amino acid exchange, excluding nonsense and frameshift variants. From that first list, we excluded the variants with the enzyme damage that has already been characterized. The clinical phenotype associated with the variants was accessed from the original article when it was not available in the databases. Variants without patient information were excluded as well as when it had been clinically associated with a nonpathogenic form of CAH or it has been found with either homozygous or compounds heterozygous with a pathogenic variant. A second screening was performed to select variants that have been found in either the Brazilian or Portuguese population.

In a second step, we analyzed the degree of amino acid conservation, chemical proprieties, and structural damage. The first two characteristics were analyzed with the software Polymorphism Phenotyping (PolyPhen-2) [31], SNAP2[32], Meta-SNP [33], PredictSNP [34], and MutPred2 [35]. To validate the methodology applied, SNPs with the known enzyme activity were used as control. Positive controls were: p.R427H (from the null group), p.I173N (from B group), and p.V282L (from C group). Negative controls were: p.R103K, p.D183E, p.M240K,and p.S269T.

For the remaining SNPs, we performed a structural impact analysis through the shift of Gibbs free energy (ΔΔG) value by STRUM [36]. Gain/loss of hydrogen bonds was calculated on WHAT-IF web (https://swift.cmbi.umcn.nl/servers/html/index.html). The human CYP21A2 structure used here was deposited on PDB by Pallan P.S., Lei L., and Egli M.[2]. This structure has residues from 28 to 485 of CYP21A2, so we added the missing amino acid with I-TASSER [37] in the N- and C-terminal. I-TASSER input was the protein sequence RefSeq NM_000500.9 and PDB ID 4y8w, chain A. The CYP21A2 structures with each variant were built with STRUM and the Gibbs free energy was compared with the wild type (WT). Structures built with the variants were aligned with the WT using PyMOL (Schrödinger, LLC) where atoms with distance variation were measured. We used the structure-naming scheme of Pallan, et al. (2015)[2].

### 4.2 Conservation analysis

To access the significance of these variants on the evolutionary conservation, we performed an evolutionary analysis with homologous proteins by ConSurf [38]. The structure of human CYP21A2 deposited under PDB ID 4y8w (chain A) was used as a reference structure and cytochrome P450 homologous sequences from *Homo sapiens, Bos taurus, Canis lupus familiaris, Cavia porcellus, Capra hircus, Felis catus, Gorilla gorilla gorilla, Lynx lynx, Mesocricetus auratus, Macaca fascicularis, Macaca mulatta, Mus musculus, Oryctolagus cuniculus, Ovis aries, Rattus norvegicus*, and *Sus scrofa* were used for conservation analysis. A multiple sequence alignment was built using CLUSTALW using Bayesian method for the calculation of conservation score. ConSurf scores range from conserved (magenta or 9) to the variable (cyan or 1). Homologous sequences were collected from SWISS-PROT with BLAST algorithm (PSI-BLAST E-value 0.0001, 4 iterations) [39-41].

### 4.3 Construction of plasmid and site-directed mutagenesis

Mammalian expression plasmid pcDNA3.1+/C-(K)-DYK carrying the ORF sequence of WT CYP21A2 (NM_000500.7) with a C-terminal DYK (FLAG) tag was purchased from GenScript (New Jersey, USA). We used this plasmid as a template to create the variants through site-directed mutagenesis. Mutagenesis reactions were performed with QuikChange II Site-Directed Mutagenesis kit (Agilent Technologies, Santa Clara, USA) following the manufacturer protocols. Oligonucleotides sequences designed with Quik Change Primer Design program (Agilent Technologies, Santa Clara, USA), are shown in **table 6**. Correct nucleotide change was confirmed by direct sequencing on ABI 3500 Genetic Analyzer (Applied Biosystems, California, USA).

**Table 6.**
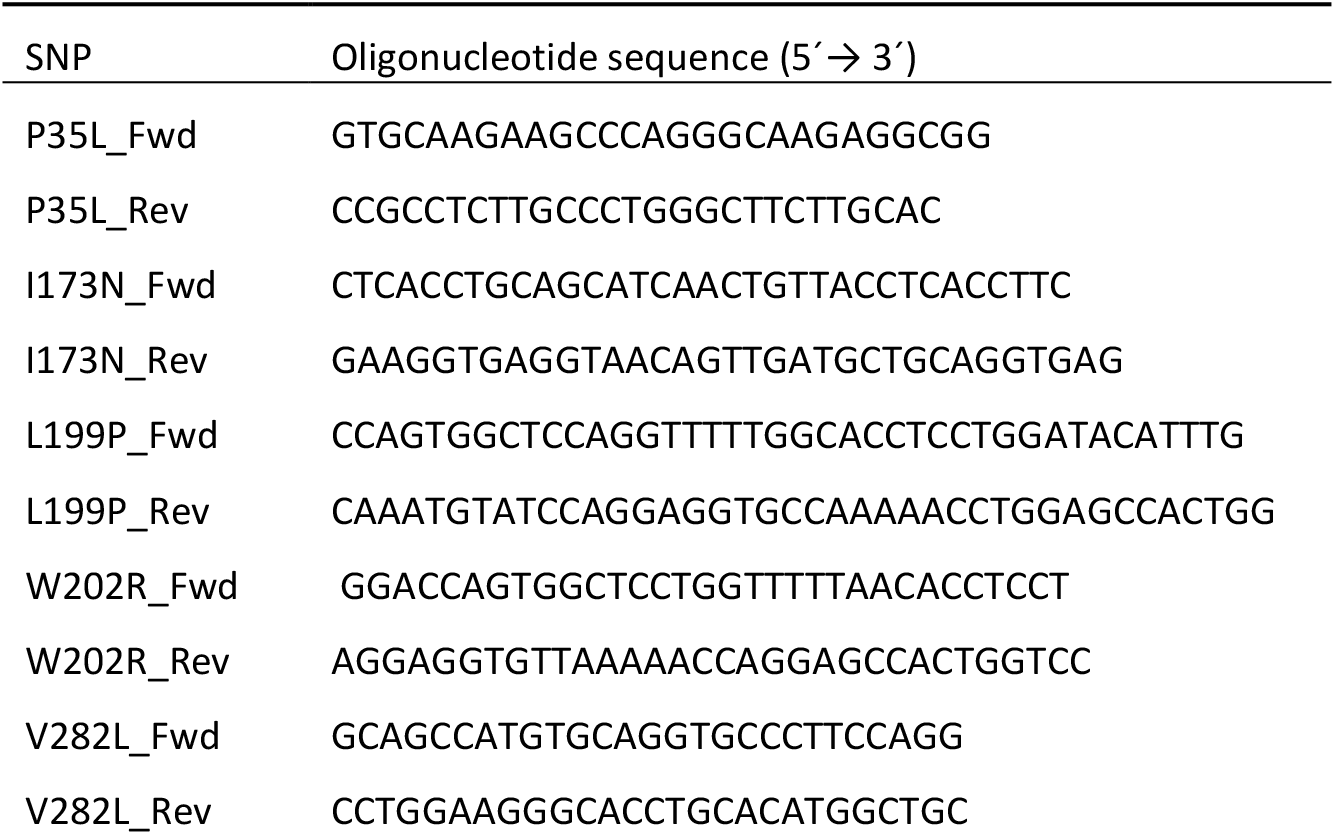
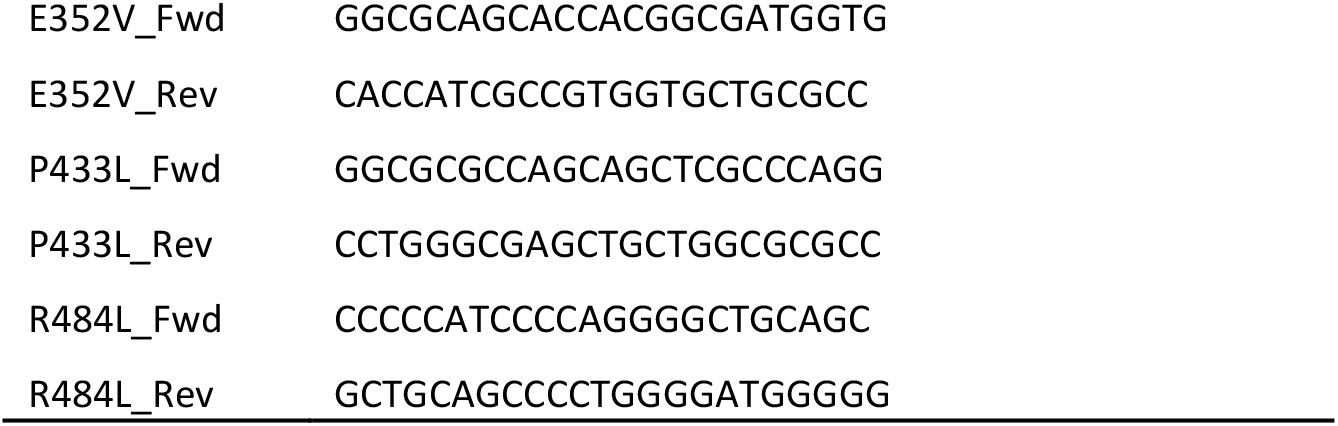
Oligonucleotides for site-direct mutagenesis. Fwd – forward, Rev – reverse.

### 4.4 Cell Transfection and enzymatic activity assay

Transient transfection was performed with the CYP21A2 WT vector and the variants. HEK293 cells were seeded in a 6-well plate (6×10^5^ cells/well). After 24 hours, growth media was replaced and the cells were transfected using Lipofectamine 2000 (Thermo Fisher, Massachusetts, USA) and 2.6 µg of the plasmid. Transfected cells were reseeded in a 24-well plate (1.5×10^5^ cells/well) after 24 hours of the transfection for uniformity of cell population across wells. At 48 h after transfection, the functional assay was started by changing to 500 µl of fresh media and adding 1 µM of unlabeled progesterone with 10,000 cpm of [^14^C]-Progesterone as a tracer. Media and cells were collected 45 min after incubation at 37 °C. Steroids were extracted from media with ethyl acetate and isooctane (1:1 vol/vol), dried, and dissolved in methylene chloride. Steroids were separated by thin-layer chromatography (TLC), exposed to a phosphor screen, and visualized with Typhoon PhosphorImager FLA-7000 (GE Healthcare Bio-Sciences AB, Uppsala, Sweden). Image intensity was measured and quantified with ImageQuant TL v8 (Cytiva, Massachusetts, USA).

The CYP21A2 enzyme activity was expressed as relative steroid conversion. Steroids were quantified as a percentage of radioactivity incorporated into 11-deoxyprogesterone to the total radioactivity measured in the whole sample and compared between the WT and variants. Cells were collected with trypsin and washed with 1x PBS to quantify the amount of protein. Results were analyzed from three technical replicates. To ensure a similar amount of CYP21A2 in each reaction, a Western blot analysis was performed to normalize the enzyme activity with the relative CYP21A2 expression.

### 4.5 Western Blot

The amount of CYP21A2 expressed was measured from the total protein extraction. Cells were incubated for 1 hour with the lysis buffer previously described [42] and centrifuged at 15,000 x g for 20 min and 4 °C. The supernatant was collected for the measurement of total protein through Pierce Coomassie Plus (Bradford) Assay Kit (Thermo Fisher, Illinois, USA). Seven µg of total protein were loaded on an SDS-PAGE gel (GenScript, New Jersey, USA), and then transferred to a PVDF membrane as previously described [42]. Two primary antibodies were used at the same time, a mouse monoclonal DKY-Tag antibody diluted 1:1,000 (GenScript, Cat# A00187) and a mouse monoclonal anti-β-Actin antibody diluted 1:1,500 (Sigma Aldrich, Missouri, USA, Cat# SAB3500350). The secondary antibody, IRDye 800CW-conjugated donkey-anti-mouse (LI-COR, Nebraska, USA - Cat# 926-32212) was diluted at 1:15,000. An Odyssey SA Infrared Imaging system (LI-COR Bioscience Inc.) was used to detect the fluorescence signal.

### 4.6 Enzyme Kinetics assay

To get the apparent reaction efficiency from the variants which showed enzyme activity, enzyme kinetic assays were performed. Five unlabeled progesterone concentrations were used (0.1, 0.3, 1, 3, and 5 µM) with 15,000 cpm of [^14^C]-Progesterone as a tracer, under the same conditions that were used in the functional assay. Enzyme velocity was normalized with the relative CYP21A2 expression derived from western blots. Results were analyzed from three biological replicates with Michaelis Menten kinetics using GraphPad Prism 9.2.0 (GraphPad, California, USA).

### 4.7 Statistics analysis

Statistical significance was calculated with a one-sample Student’s *t*-test, for comparing samples in the same group, and one-way ANOVA, for comparing two groups (one variant group with the WT group). The P-value was considered significant with p < 0.05. Prism 8 (GraphPad, California, USA) and Excel (Microsoft, Washington, USA) were used to perform the calculation and statistical analysis.

## 5. Conclusions

Here we elucidated the structural and functional impact of six CYP21A2 variants (p.P35L, p.L199P, p.W202R, p.E352V, p.P433L, and p.R484L) with unknown profiles via *in silico* and *in vitro* enzyme analysis. We showed a good correlation between *in silico* and *in vitro* functional studies for structural and conserved variants. However, residues with specific roles have ambiguous results, e.g. for the catalytic site, as the homology has a high weight for most of the predictor tools. Especially in these cases, the functional assay to access the enzyme activity was decisive to assign the damage caused by that variant. These results also emphasize the relevance of using multiple algorithms together with functional assays especially when it is not possible to establish correlations with the phenotype-based solely on bioinformatics predictions.

## Supplementary Materials

The following are available online at www.mdpi.com/xxx/s1, Figure S1: Specific 21-hydroxylase (21OH) protein expression by western blot, Figure S2: Thin-layer chromatography (TLC) from the kinetics assays with P35L, L199P and P433L, Figure S3: Plot of the 21-hydroxylase specific activity, Table S1: Hydrogen-bounds of each wide-type residues were studied with the CYP21A2 structure, Table S2: Hydrogen-bounds of the variants studied on the CYP21A2.

## Author Contributions

Conceptualization, M.J.P. and A.Z., M.L.R.R., A.V.P; methodology, M.J.P., A.V.P, R.L.B., S.S., B.V.M., and T.R.; formal analysis, M.J.P., and A.V.P.; investigation, M.J.P.; resources, A.V.P. and A.Z.; writing—original draft preparation, M.J.P; writing—review and editing, A.Z., and A.V.P.; visualization, M.J.P and A.V.P.; supervision, A.V.P., A.Z., M.L.R.R., and R.L.B.; project administration, A.V.P.; funding acquisition, M.J.P., A.V.P., A.Z., and K.M. All authors have read and agreed to the published version of the manuscript.

## Acknowledgments

The authors are grateful to Christa E. Flück, Kay Sauter, and Maria Natalia Rojas Velazquez for help with the experimental setup, enzyme assays and valuable discussions.

