## Supplemental Data for "Characterization of mutations causing steroid 21-hydroxylase deficiency in Brazilian and Portuguese populations"

[1] Graduate Program in Cell and Molecular Biology, Universidade Federal do Rio Grande do Sul (UFRGS), Porto Alegre CEP 91501-970, Brazil.

[2] Center for Biotechnology, Universidade Federal do Rio Grande do Sul (UFRGS), Porto Alegre CEP 91501-970, RS, Brazil.

[3] Department of Biomedical Research, University of Bern, Bern 3010, Switzerland.

[4] Pediatric Endocrinology Unit, Department of Pediatrics, University Children's Hospital Bern, Bern 3010, Switzerland

[5] Departament of Pharmacosciences, Universidade Federal de Ciências da Saúde de Porto Alegre (UFCSPA), Porto Alegre CEP 90050-170, Brazil.

[6] Serviço de Referência em Triagem Neonatal, Hospital Materno Infantil Presidente Vargas, Porto Alegre CEP 90035-074, Brazil.

[7] Graduate Program in Molecular Biology Applied to Health, Universiade Luterana do Brasil (ULBRA), Canoas CEP 92425-020, Brazil.

‡These authors contributed equally to this work.

(A)

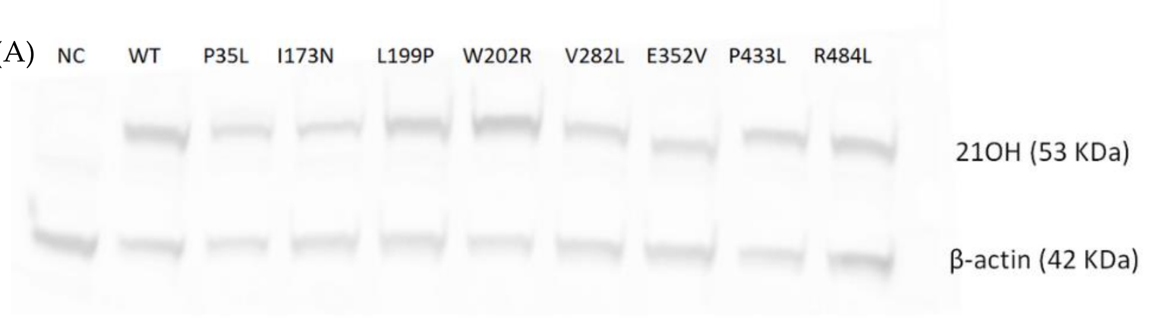


(B)

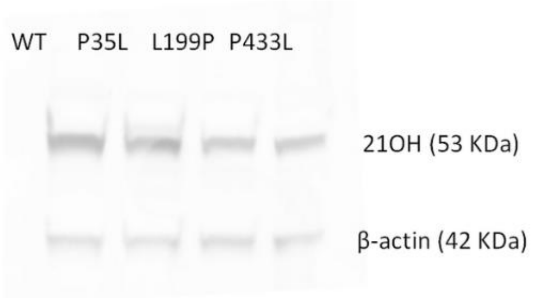


**Figure S1.** Specific 21-hydroxylase (21OH) protein expression by western blot. Cells from the functional and kinetic assay were collected and lysed. The total protein was measured through Pierce Coomassie Plus (Bradford) Assay Kit. Seven μg of total protein were loaded on an SDS-PAGE gel and then transferred to a PVDF membrane. Two primary antibodies were used, a mouse monoclonal anti-flag for the 21OH (53 KDa) and a mouse monoclonal anti-β-actin for the normalizing gene β-actin (42 KDa). One secondary antibody was used, IRDye 800CW-conjugated donkey-anti-mouse. The fluorescence signal was detected with Odyssey SA Infrared Imaging system. (A) 21OH protein from the functional assay with all variants. Western blot was performed for one sample of the technical triplicate. (B) 21OH protein from the kinetic assay. Western blot was performed for one sample of the biological triplicate. NC, negative control (basal HEK293 cells); WT, 21OH wild type.

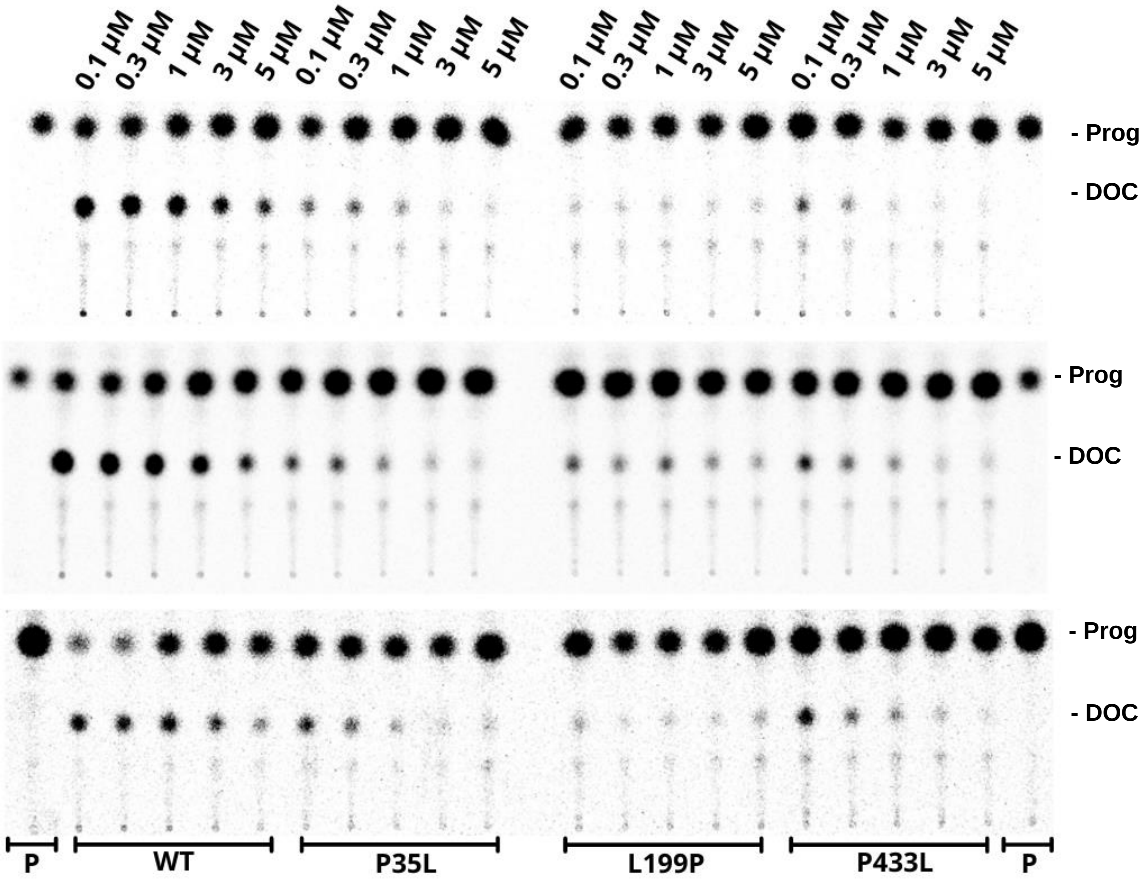


**Figure S2.** Thin layer chromatography (TLC) from the kinetics assays with P35L, L199P and P433L. The 21-hydroxylase activity was measured by the conversion of progesterone (Prog; P) (0.1, 0.3, 1, 3 and 5 μM) with 15,000 cpm of [$^{14}$C]-Progesterone into 11-deoxycorticosterone (DOC) radioactive. Three biological replicates were performed. WT, wild type.

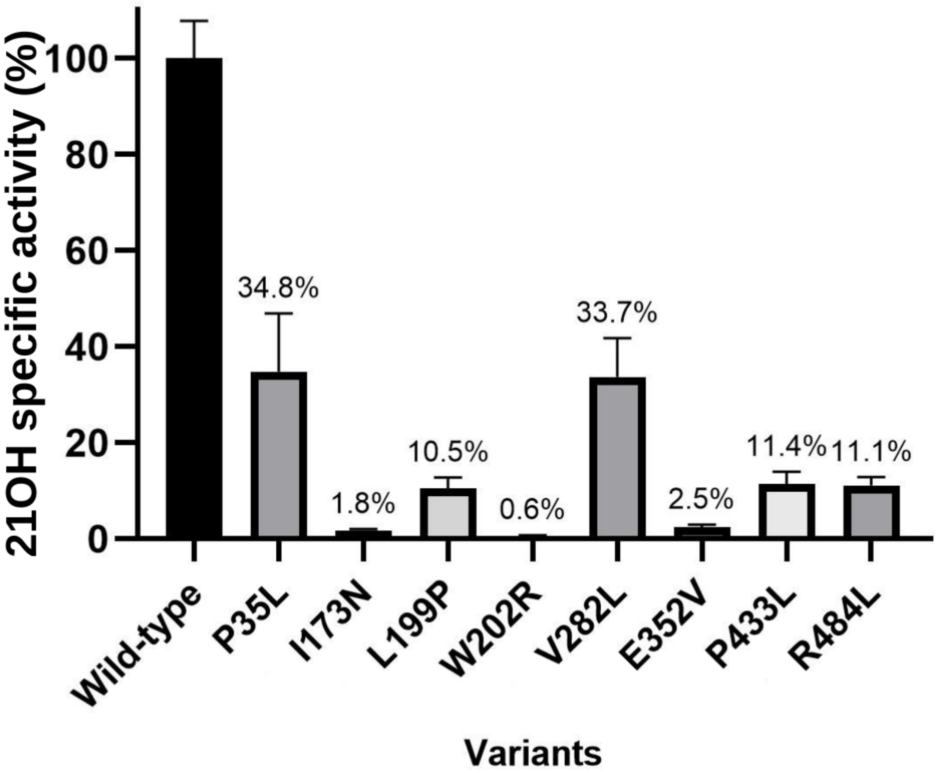


**Figure S3.** Plot of the 21-hydroxylase specific activity. The specific activities are expressed by the enzyme activity obtained from the TLC spot densitometry by the specific protein estimated through western blot. Results are from a technical triplicate, and all calculations were done on GraphPad software.

**Table S1.** Hydrogen-bounds of each wide-type residues studied on the CYP21A2 structure. The CYP21A2 structure was built on I-TASSER based on the x-ray crystallography structure deposited on PDB under the ID 4Y8W. Hydrogen bonds were calculated with the Optimal Hydrogen Bounding Network tool on WHAT-IF web (https://swift.cmbi.umcn.nl/servers/html/index.html).

| Amino acid 1 | Atom 1 | Amino acid 2 | Atom 2 | Distance 1-2 (Å) |
|---|---|---|---|---|
| LEU 199 | N | ILE 195 | O | 3.11 |
| LEU 199 | O | SER 203 | N | 3.00 |
| TRP 202 | N | VAL 198 | O | 3.00 |
| GLU 352 | N | ALA 348 | O | 2.99 |
| GLU 352 | OE2 | ARG 355 | NE | 2.80 |
| GLU 352 | OE1 | ARG 355 | NH1 | 2.78 |
| GLU 352 | O | LEU 356 | N | 2.97 |
| GLU 352 | OE1 | TRP 406 | N | 2.86 |
| PRO 433 | O | ALA 435 | N | 2.64 |
| ARG 484 | N | GLN 482 | O | 3.07 |
| ARG 484 | N | ALA 449 | O | 2.98 |
| ARG 484 | O | MET 486 | N | 2.96 |

**Table S2.** Hydrogen-bounds of the variants studied on the CYP21A2. The complete CYP21A2 protein structure based on the PDB ID 4Y8W was mutated on STRUM (one mutation per structure). Hydrogen bonds were calculated with the Optimal Hydrogen Bounding Network tool on WHAT-IF web (https://swift.cmbi.umcn.nl/servers/html/index.html).

| Amino acid 1 | Atom 1 | Amino acid 2 | Atom 2 | Distance 1-2 (Å) |
|---|---|---|---|---|
| PRO 199 | O | SER 203 | N | 2.79 |
| ARG 202 | N | VAL 198 | O | 3.02 |
| VAL 352 | N | ALA 348 | O | 2.97 |
| VAL 352 | O | LEU 356 | N | 2.91 |
| LEU 433 | O | ALA 435 | N | 3.16 |
| LEU 484 | N | ALA 449 | O | 2.85 |
